# Korean Natural Farming practices are dominated by a limited number of microbes and decrease fungal diversity

**DOI:** 10.1101/2025.09.26.678907

**Authors:** Candace Thompson, Shawn Mozeika, Elizabeth Paredes, UnJin Lee

## Abstract

Korean Natural Farming (KNF) practices claim to cultivate and transfer “indigenous micro-organisms” (IMOs) to donor soils as a method of probiotic soil enhancement. We investigated whether IMO cultivation can propagate unique microbiomes and maintain microbial diversity through successive IMO stages for restoration of flood contaminated soils. Employing a balanced study design using soil samples from salt marsh, deciduous forest, and urban greenspace (plus sterilized controls), samples underwent the first two IMO cultivation steps followed by 16S rRNA and ITS metagenomic sequencing. Notably, IMO cultivation was dominated by limited bacterial taxa (*Enterobacterales*, *Pseudomonadales*, *Bacillales*) and fungal taxa (*Rhizopodaceae*, particularly *R. oryzae*). While bacterial diversity was maintained or increased during two IMO stages, fungal diversity consistently decreased. Principal Coordinates Analysis also revealed distinct clustering by inoculum source (i.e. human-altered, human-transported (HAHT) vs. natural vs. sterile) that persisted throughout cultivation. Our evidence suggests that the IMO process enriches for specific taxa likely adapted to cultivated conditions and fails to maintain fungal diversity, contrasting greatly with KNF’s proposed benefit of propagating locale-specific, fungal-dominated indigenous microbiomes. However, our results demonstrate that early IMO cultures may capture and sustain bacterial diversity in soil, opening the door for future studies of KNF efficacy and sustainability.

**Sustainability Statement:** This work is primarily associated with UN Sustainable Development Goal SDG 15 (Life on Land). By scientifically evaluating Korean Natural Farming “indigenous Microorganism” (IMO) practices through metagenomic sequencing, this study provides evidence-based insights into microbial community dynamics relevant to soil restoration and terrestrial ecosystem health. The findings inform low-cost bioremediation strategies for flood- contaminated urban soils, which is of growing importance as climate-driven extreme precipitation events intensify. Additional relevant SDGs include: SDG 13 (Climate Action), SDG 11 (Sustainable Cities and Communities), SDG 2 (Zero Hunger), and SDG 3 (Good Health and Well-being).

## Introduction

Healthy soils underpin human and ecological well-being through food production, nutrient cycling, carbon sequestration, and the support of global biodiversity (Wagg *et al*. 2021, Telo da Gama 2023). Yet soils worldwide are becoming increasingly depleted due to the accelerating impacts of the climate crisis (Qu *et al*. 2024, Kopittke *et al*. 2025). Climate-driven high-volume rain events and subsequent flooding impact soils by reducing soil oxygen and spreading pathogens, heavy metals, and other contaminants (Unger, Kennedy, and Muzika 2009, Shi *et al*. 2010, Quist *et al*. 2022, Furtak and Wolińska 2023, Seneviratne *et al*., n.d.). From storms depositing 1–3 inches of rain per hour to major hurricanes such as Ida and Sandy (Rosenzweig and Solecki 2014, Fowler *et al*. 2021, Mossel *et al*. 2024), New York City has been subjected to a similar recent increase in climate-induced flash flood events. Such events pose both immediate and long-term risks (Kovats *et al*. 2024), with projections indicating that approximately 30% of the city’s land area will be flood-prone by 2080 (Patrick *et al*. 2015). Finding low cost and accessible methods for restoring healthy soil ecology is of increasing importance and pertains to the resilience of agricultural systems, habitat conservation, and land stewardship.

Stuyvesant Cove Park (SCP) is a two-acre, all-native plant park located in lower Manhattan that was recently razed and reconstructed as part of the “The Big U”: a citywide climate mitigation project that seeks to protect lower Manhattan through an interconnected chain of gray infrastructure, floodwalls, and berms. Extending from 23rd to 18th Streets, “Stuy Cove” is situated between the East River Estuary and a newly constructed floodwall that spans 5 city blocks. During future storm events the site is designed to absorb tidal impacts, temporarily inundating the landscape while protecting the critical utilities, businesses, and residences that lie on the western “dry” side of the wall. However, this tract of land was once the site of a manufactured gas plant which has since been capped and remediated, with contaminated subsoils and containment wells that regularly test positive for high levels of coal tar, benzene, toluene, ethylbenzene, xylene, and more complex PAHs (NYSDEC, and conEdison 2017a, 2017b). During storm events these contaminants are likely to mobilize and spread across the clean soils of this nascent landscape. Stuyvesant Cove Park thus presents unique opportunities as a living laboratory, allowing for the exploration of affordable bioremediation strategies that can potentially restore soils to healthy baselines following future flood events.

One promising technique popular within the regenerative agriculture community is the Korean Natural Farming (KNF) practice known as “indigenous Microorganisms” (IMO). This low-tech process involves collecting microbial communities from intact ecosystems—such as old-growth forests or salt marshes—and culturing them through a series of different substrates, e.g. rice (“IMO1”), unrefined sugars/jagggery (“IMO2”), rice bran/flour (“IMO3”), field soil/red fine soil (“IMO4”), to create a probiotic soil amendment that can be applied to degraded or unhealthy soils (Park and DuPonte 2008, Cho and Cho 2010, Reddy 2011, Keliikuli 2018, Keliikuli *et al*. 2019).

Despite increasing interest and claims of efficacy among regenerative agriculture practitioners, peer reviewed research on this practice remains limited. Instead, the majority of the extant literature on the topic remains primary sources (Cho and Cho 2010, Reddy 2011) or are technical documents that have not been peer review (Park and DuPonte 2008, Keliikuli 2018, Keliikuli *et al*. 2019). While this paucity of work makes it difficult to identify proposed mechanisms of IMO function, it could potentially involve the cultivation of key microbes that either increase the availability of critical nutrients in a manner similar to fertilizer (Keliikuli *et al*. 2019), or increase plant growth through the presence of plant growth-promoting rhizobacteria (PGPR)(Keliikuli *et al*. 2019). However, the name suggests that some intrinsic quality of these naturally occurring microbes’ indigeneity, as opposed to a specific set of of site-independent microbes, is a key factor in the function of the “indigenous micro-organism” process. Questions regarding the microbial dynamics of Korean Natural Farming practices thus remain unresolved, including which, if any, microbes recur in the IMO process independently of soil inoculant source, and what degree the IMO process can capture and/or propagate diversity.

To test the microbial dynamics of the KNF process, we selected two ecologically intact “natural” sites to serve as potential donor cultures for Stuyvesant Cove Park: (1) Marshlands Conservancy, one of the few remaining saltwater marshes in the greater New York area, adapted to daily brackish inundation and representative of Stuy Cove’s preindustrial ecosystem; and (2) the Thain Family Forest at the New York Botanical Garden, one of the last remaining old-growth forest fragments in the region and therefore likely a fungally-dominant soil ecology (Byers *et al*. 2020, Hu *et al*. 2024). While these “untouched” or “wild” northeastern landscapes do likely host endemic and long existing soil biota, they are also sited within one of the most densely populated metropolitan areas in America, and equally subjected to common anthropogenic stressors such as petrochemicals, heavy metals, raw sewage, and more (Byers *et al*. 2020). We therefore hypothesized that these legacy soil horizons could harbor both diverse and resilient communities capable of serving as robust donor microbiomes for the newly installed human-altered, human-transported (HAHT) soils at Stuyvesant Cove (see Methods and Materials). Specifically, we assessed whether native microbial communities from each soil source are able to be propagated throughout the IMO process, thus allowing for later usage as donor microbiomes.

We report findings from the early stages of the IMO process (i.e. soil, IMO1, and IMO2 stages), using a replicable and balanced study design to examine the IMO process’ underlying microbial dynamics via 16S and ITS metagenomic sequencing. For this initial study we executed the first two stages of the IMO process in triplicate at each donor site and at Stuyvesant Cove Park, utilizing soil samples, IMO cultures, and sterilized controls. We extracted environmental DNA (eDNA) from these biological replicates, amplifying and sequencing 16S and ITS metagenomic libraries from multiple intermediate time points. We examined unweighted within- and between-group β-diversity dynamics of these cultures and reported changes in key organized taxonomic units (OTUs). Our pilot study thus addresses several strong hypotheses about the IMO process: to what extent sequential selection events using differing carbon sources alter microbiome composition, to what extent inoculants determine the dynamics of microbial community succession, whether community succession in IMO cultivation is robust to initial conditions, and finally, whether microbial diversity can be maintained and propagated using IMO media.

## Materials and Methods

### Site Selection and Sample Collection

Soil samples were collected from three ecological sites in the New York metropolitan area. Extant salt marsh samples were obtained with permission from the Marshlands Conservancy (“marshlands”) in Rye, New York. Temperate deciduous forest samples were collected with permission from the New York Botanical Garden’s Thain Family Forest (“forest”), one of the few remaining old-growth forests in the city. Soils from Stuyvesant Cove Park, a recently reconstructed park were also collected. Importantly, the soils from Stuyvesant Cove Park are urbanogenic human-altered, human-transported (HAHT) material (Soil Survey Staff 2022), consisting of a naturally-sourced topsoil amended with biochar and mulch top dressing. This HAHT material was then placed atop a gravel drainage later and was subject to high human traffic over a period of approximately one year.

To control for potential weather-related externalities, all sample collection and IMO installations were performed on September 16^th^, 2024 for all sites. Similarly, completed IMO1 cultures were retrieved and processed on September 20^th^, 2024. No precipitation occurred during the collection and incubation period, while previous recorded rainfall occurred on September 8^th^, 2024 at 0.2 inches total.

At each location, three test plots were selected in collaboration with local staff and sited within 20 paces of each other. The Marshlands Conservancy plots were located at 40°56’52.3“N 73°42’02.8“W, New York Botanical Garden plots were located at 40°52’02.2“N 73°52’31.9“W, and Stuyvesant Cove Park plots were located at 40°43’57.6“N 73°58’25.7“W, Plots were selected for their slight but notable differences in terrain or slope, tide proximity, sun exposure, and adjacent plant or fungal communities. Ambient temperature and humidity, and other observations during collections are provided in Table S1. Marshlands Conservancy plots are classified as Charlton-Chatfield complex (CrC), New York Botanical Gardens plots are classified as Fluventic Hapludolls-Cumulic Endoaqolls complex (FFA), and Stuyvesant Cove Park plots are classified as Urban lan-Laguardia complex (ULAI) according to the USDA Web Soil Survey (Soil Survey Staff 2026).

500g of soil was collected from the top 10-20 cm of the soil horizon in each test plot after removal of surface humus. Depth was measured using a demarcated hori knife. Prior to each collection, digging tools were field sterilized using 70% ethanol. Each soil sample was placed into a sterile, plastic bag which was vigorously shaken until completely homogenized. One-gram subsamples were extracted from the homogenized 500g sample using ethanol-sterilized tools and subsequently suspended in 10 mL of 0.9% NaCl. The resulting solution was vigorously shaken by hand for 1 minute in the field. 200 µL of the suspended solution was immediately transferred into Zymo DNA/RNA Shield Lysis and Collection Tubes (R1103) using Eppendorf micropipettes and fresh, sterile pipette tips.

The remaining ∼500g of bulk dry soil samples from each test plot was then compiled into a master composite site sample into a new, sterile bag. The resulting soil mix was homogenized again and returned to the lab. Later that day, 1 gram samples were collected from each composite site sample using ethanol-sterilized tools and immediately shipped overnight for chemical and biological characterization by Logan Labs (member of the North American Proficiency Testing Program) and Rhizos LLC (Table S2). Fresh, sterile nitrile gloves were worn to ensure proper sterile technique throughout. All soil sample collection and processing steps were performed by CT and EP in order to minimize batch effects.

### IMO Cultivation

The IMO1 and IMO2 collection process was carried out in accordance with (Cho and Cho 2010) and (Reddy 2011). IMO1 collection boxes were constructed from cedar 1 x 4 in. planks (12 × 12 × 4 in.) covered in galvanized steel mesh (Figure S1). Slightly undercooked, sterilized organic brown rice (300 g) was added, the boxes were screwed shut and placed into plastic bags for transport to their install site. While (Reddy 2011) state that white or brown rice may be used interchangeably, brown rice was selected as it was deemed to be a more nutritious substrate for fungi cultivation.

Once on site, the surface humus was removed and the IMO box was installed where the rice could make full contact with the soil horizon through the hardware cloth mesh. A piece of nonwoven Remay**©** row cover was placed over the top of the box, and the removed surface humus was laid over top the cloth as a “soil sandwich.” A piece of cardboard was then tented over the box and staked with landscape pins to allow airflow while also providing shelter from potential rain storms. These boxes remained installed for four days, during which there was no notable rainfall. Upon retrieval, surface humus and row cover were removed and the box was slowly lifted from the soil surface and placed into a sterilized bag, taped shut, and transported to our lab for further processing (Figure S1). All installations were performed on the same days by CT and EP in order to minimize batch effects.

After collection, IMO1 cultures were unboxed in their entirety on the same day, weighed, placed into a sterilized glass bowl, and homogenized by hand using sterile, nitrile gloves. 1g of substrate was collected from various points in the larger sample using sterilized tools and resuspended in 10 mL of 0.9% NaCl, which was vigorously shaken form 1 minute. From this, 200 µL was stored in Zymo DNA/RNA Shield Lysis and Collection Tubes. Mycelial growth, typically white and down-like, was observed, and pure samples were isolated for library extraction (Figure S1c,e). Two of the “forest” samples were tampered with by wildlife (likely raccoons), failed to culture, and therefore were discarded. Therefore, the 3 “marshlands” and remaining “forest” sample were pooled as “natural” samples for comparison with Stuyvesant Cove’s HAHT soils (see <u>Experimental design enables differentiation between HAHT, natural, and control microbiomes</u>).

For IMO2 production, homogenized IMO1 material was mixed with jaggery sugar at a 1:1 ratio (w/w) and incubated at room temperature under aerobic conditions for seven days. Jaggery sugar was selected in accordance with (Reddy 2011), being the least refined choice between jaggery, brown sugar, and white sugar. These samples were contained within sanitized mason jars covered with several layers of clean Remay**©**; an aerobic environment (Figure S1f). Samples were collected on days 3 and 7, processed as above, and stored at 4°C until DNA extraction. All IMO2 material was cultured in the same room on the same days and processed solely by CT in order to minimize potential batch effects.

As mentioned previously, 500g soil samples from each test plot were collected prior to IMO box installation and removal. The three soil samples from each location were combined into an aggregate sample, and sterilization controls were prepared by autoclaving at 121°C for 30 min. These soils were then installed in sanitized plastic bins drilled with ½” aeration holes covered by 3M micropore surgical tape at a 2” spacing. (Figure S1f), which is an imitation of the common monotub TEK used for at-home mushroom cultivation by the DIY mycology community. New IMO boxes filled with organic brown rice were placed in these tubes, the rice making full contact with the soil surface, and were left to culture in an open air room at ambient air temperature out of direct sunlight for 4 days. These cultures were generated synchronously by CT with other IMO1 cultures in order to reduce potential batch effects.

### DNA Extraction and Sequencing

All samples were stored in Zymo DNA/RNA Shield Lysis and Collection Tubes containing Zymo Bashing Beads. Tubes were brought to room temperature, vortexed using a Scientific Industries Multi-tube Holder (SIH524), and total environmental DNA was extracted using ZymoBIOMICS DNA Miniprep Kits (D4304). DNA concentrations were quantified by Nanodrop (Thermo Scientific NanoDrop OneC). All libraries extracted on the same day by UL to minimize batch effects and were subsequently blinded by randomly assigning library labels. Unblinding was performed only during final analysis steps.

Amplicon libraries were generated using Zymo Quick-16S Plus NGS Library Prep Kits (V4; D6430) and Quick-ITS Plus NGS Library Prep Kits (UDI; D6425) according to manufacturer protocols. Amplifications were compared against positive (ZymoBIOMICS Microbial Community Standards) and negative (ddH O) controls. Libraries with poor amplification were processed using Zymo Oligo Clean & Concentrator Kits (11-380B) prior to re-amplification. Notably, all 16S libraries were prepared simultaneously on the same day by UL, while all ITS libraries were prepared simultaneously on another day by UL in order to minimize batch effects. Libraries were pooled in equal volumes according to manufacturer protocol, cleaned, and prepared with a 15% PhiX spike-in. Sequencing was performed by the Rockefeller University Genomics Resource Center (RRID: SCR_020986) on an Element Biosciences AVITI platform (2 × 300 bp, Cloudbreak Freestyle kit, 300M read depth) in a single, exclusive run in order to minimize batch effects. From 84 libraries submitted, yields ranged from 183.7 Mb to 5.47 Gb. Sequencing metrics and index assignments are provided in Table S3.

### Bioinformatic Analysis

Raw reads were processed using QIIME2 (version 2024.10, quay.io distribution)(Bolyen *et al*. 2019). Demultiplexing and quality filtering were performed with the q2-demux and q2-dada2 plugins. Taxonomic assignment employed the SILVA database (release 138)(Glöckner *et al*. 2017) for 16S sequences and the UNITE database (version 10)(Abarenkov *et al*. 2024) for ITS sequences. Sequences lacking phylum-level classification were removed prior to analysis, while sequences that were incorrectly mapped were also removed (i.e. 16S sequences matching eukaryotic clades, or ITS sequences matching non-eukaryotic clades). Additionally, sequences mapping to mitochondrial sequences were also removed prior to analysis. All datasets were blinded before analysis.

Diversity metrics including unweighted UniFrac (Lozupone *et al*. 2011) and Jaccard indices (Mainali *et al*. 2017) were calculated at a rarefaction depth of 280,000 reads, chosen to match the smallest quality-filtered library. Differential abundance testing was performed using ANCOM-BC (Lin and Peddada 2020) in QIIME2 with a significance threshold of p = 0.001. α- and β-diversity metrics were calculated in QIIME2. Functional predictions were generated using the PICRUst2 (Douglas *et al*. 2020) plugin (‘q2-picrust2’, v2024.5) for QIIME2 contained in the EBI Mgnify (Richardson *et al*. 2022) pipeline (microbiome-informatics/q2-picrust2 distribution on quay.io). Differential abundance calculations for pathway enrichment were performed using LinDA (p < 0.05 with Benjamini-Hochberg correction)(Zhou *et al*. 2022) and visualized using ggpicrust2(Yang *et al*. 2023). Relative abundance and UniFrac distributions were analyzed and visualized in R. Scripts are available at https://github.com/ulee-sciscripts/imo

## Results

### Experimental design enables differentiation between HAHT, natural, and control microbiomes

To test whether inoculation source altered microbial community dynamics, we employed a balanced experimental design utilizing soils collected from natural environments and a HAHT greenspace. Physical characteristics of soil specimens are reported in Table S2 and α-diversity metrics are reported in Figure S2. IMO1 cultures were inoculated *in situ* at each source and compared with IMO1 cultures initiated from sterile controls (Methods and Materials, Figure 1). Since IMO1 cultures require low-moisture, aerobic conditions, our sterilized control IMO1 cultures may potentially have been inoculated via airborne particulates or aerosols, rather than residual soil inocula (Methods and Materials). As all IMO1 cultures were also subjected to similar levels of airflow, the sterile controls may potentially reflect effects of passive inoculation via the experimental environment. Sterilized control IMO1 samples were cultured under shared laboratory conditions to yield IMO2 cultures. To ensure replicability, biological triplicates were generated from each *in situ* soil source, as well as one sterilized control per location. The following samples were collected at each stage of the IMO process: live soil (with one autoclaved control), IMO1 (field and laboratory replicates), and IMO2 at days 3 and 7. Two forest replicates were lost due to wildlife disturbance; therefore, forest and marshland samples were pooled as “natural,” Stuyvesant Cove samples as HAHT, and autoclaved soils as “sterile”. The decision to pool forest and marshland samples is supported by the observation that the single forest and all three marsh samples cluster more closely than sterilized and HAHT samples across the first three principal components in both 16S and ITS data (Figure S3a-d). This remains true in soil samples and in IMO samples, thus demonstrating that the effect of forest vs marsh origin is smaller than the effect of soil, IMO, and staging effects. Additionally, pairwise Unweighted Unifrac distance calculations demonstrate that the distribution of distances between different marshlands samples is similar to distances between marshlands and forest samples (Figure S4, S5)(see <u>Diversity dynamics during cultivation</u>). Such classification is also consistent with both marshland and forest sites being designated as flood-tolerant soil types according to the USDA Soil Survey (Soil Survey Staff 2026). This balanced experimental design thus allowed us to identify specific inoculation effects while controlling for environmental effects.

**Figure 1.**
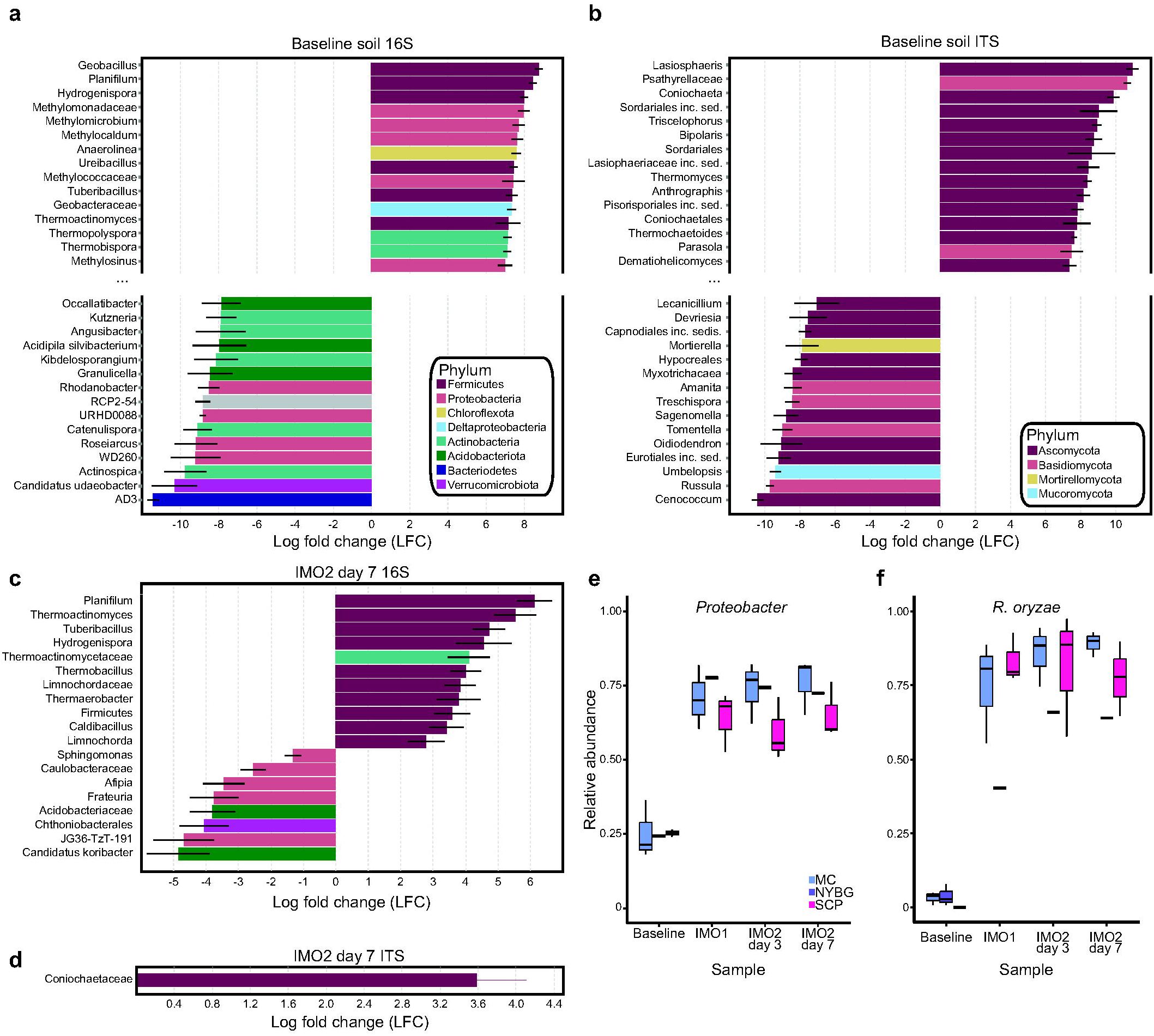
IMO process and study design. The IMO process begins by *in situ* inoculation of a carbohydrate source (sterilized rice) using live soil samples producing IMO1. The IMO1 is then homogenized then transferred onto a second carbohydrate source (jaggery) and allowed to ferment producing IMO2. This is then combined with a third carbohydrate source (rice bran) to produce IMO3, which is then mixed with soil to produce the soil amendment IMO4. The study design utilized three soil sources as inoculants for IMO1. Soil was then sterilized and also used as an IMO1 starter. IMO2 was then cultured in shared environmental conditions and subsequently sampled at day 3 and day 7 Alt text: Flowchart diagram detailing the four-stage Indigenous Microorganism (IMO) workflow alongside a regional map and microbial diversity trends. On the left, a map highlights sampling locations in New York: Marshlands Conservatory, New York Botanical Garden, and Stuyvesant Cove Park. The central flowchart visualizes the physical steps from soil samples to buckets and vials representing IMO1, IMO2, IMO3, and IMO4, including a parallel sterile control path. DNA icons mark specific library prep collection points throughout the stages. A graph at the bottom indicates that bacterial diversity increases while fungal diversity decreases from Stage 1 to Stage 4.

### Taxonomic dynamics during cultivation

To get an overview of the IMO process, we first examined the relative abundance of key species, identified significantly differentially abundant taxa, and subsequently characterized their dynamics (Figure 2). To determine whether our experimental strategy produced sufficiently accurate results, we performed a taxonomic analysis of our 16S and ITS libraries amplified from the synthetic Zymo Community Standard. Both 16S and ITS taxonomic analyses demonstrated high accuracy at our down-sampled depth when compared to the Zymo Community Standard. For 16S communities, observed abundances closely matched expected values (*P. aeruginosa*: 4.2% expected vs. 3.5% observed, *E. coli*: 10.1% expected vs. 11.6% observed, *S. enterica*: 10.4% expected vs. 10.8% observed, *L. fermentum*: 18.4% expected vs. 16.8% observed, *E. faecalis*: 9.9% expected, 9.1% observed, *S. aureus*: 15.5% expected vs. 12.9% observed, *L. monocytogenes*: 14.1% expected vs 13.1% observed, *B. subtilis:* 17.4% expected, 22.1% observed). For ITS communities, *S. cerevisiae* and *C. neoformans* were detected at expected ratios (50% expected vs. 50.1% observed and 50% expected vs. 49.9% observed, respectively).

**Figure 2.**
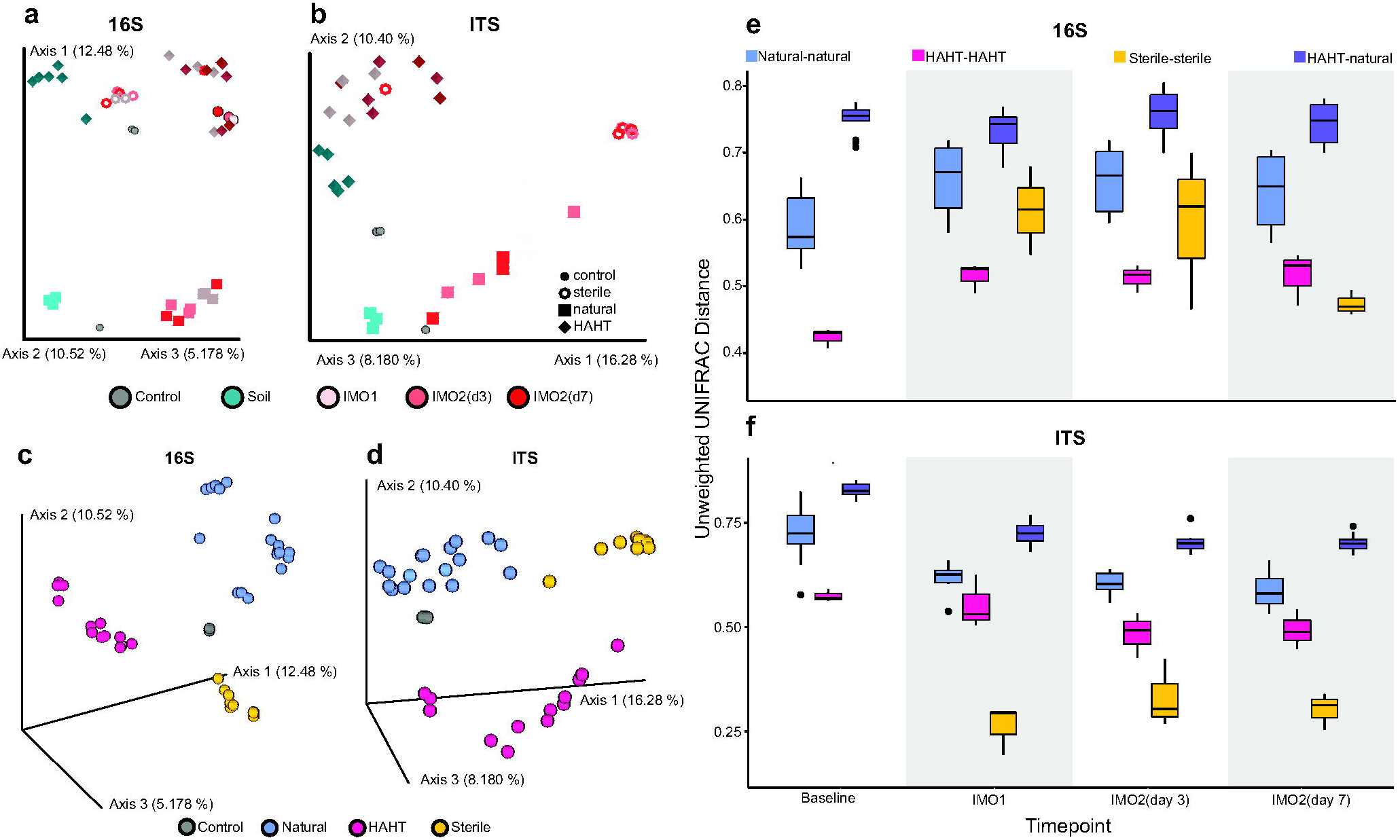
Taxonomic analysis of IMO process reveals that later IMO stages are dominated by limited taxa. Relative abundances of top 15-most prevalent organized taxonomic units (OTUs) for (a) 16S or (b) ITS libraries. Libraries are labeled by location, IMO stage, and treatment status. Libraries are grouped left to right: synthetic or otherwise special communities (EM1 = commercially available product, COMM = Zymo community standard, Mycellium = mycellium sample, see text), sterilized samples, HAHT-sourced samples, and natural samples. Locations: MC = Marshland Conservancy, NYBG = New York Botanical Garden, SCP = Stuyvesant Cove Park. Alt text: Two stacked bar charts, labeled a and b, showing taxonomic relative frequency from 0% to 100% across various samples and experimental stages. Chart a represents 16S bacterial libraries, and chart b represents ITS fungal libraries. The x-axis groups vertical bars into categories: unique controls, sterile samples, HAHT samples, and natural samples, subdivided by stage from baseline to IMO2 day 7. Legends on the right color-code the top 15 taxonomic families. Visually, earlier stages show a diverse multicolored breakdown, while the later IMO2 day 3 and day 7 stages become dominated by single, solid blocks of color, representing a sharp drop in overall community diversity.

Relative microbe abundance obtained from 16S and ITS sequencing suggests an immediate and striking difference between untreated soil and IMO microbiomes. Microbiomes derived from HAHT (i.e. Stuyvesant Cove Park) and “natural” (i.e. Marshland Conservancy and Thain Family Forest, see **PCoA Reveals IMO Staging and Inoculant Source Effects on Bacterial and Fungal Composition**) soil samples show that the vast majority of their microbial communities lie outside of the top 15 most abundant Organized Taxonomic Units (OTUs). Once the IMO process began, community composition shifted rapidly toward dominance by a small number of bacterial (i.e. *Enterobacterales*, *Pseudomonadales*, *Bacillales*) and fungal (i.e. *Rhizopodaceae*) clades. The majority of microbial composition in nearly every IMO sample consisted of these clades, while the proportion of other taxa decreased over successive stages (Figure 2).

“Sterile” cultures’ microbial compositions were dominated by *Bacillales* (e.g. *Bacillus subtilis*), HAHT cultures by *Enterobacterales* (e.g. *Escherichia coli*) and *Bacillales*, and natural cultures by *Enterobacterales*, *Pseudomonadales* (e.g. *Pseudomonas aeruginosa*), and *Burkholderiales*. All IMO cultures, regardless of source, also showed a high prevalence of *Rhizopus oryzae* (*Rhizopodaceae*), a filamentous fungus used in traditional tempeh fermentation and industrial food production (Figure 2b). This observation is consistent with visual observation of such mycelia across all IMO samples regardless of origin or sterilization treatment (Figure S1c,e).

An analysis of compositions of microbiomes with bias correction (ANCOM-BC, p < 0.001), which identifies differentially abundant taxa while accounting for absolute abundance and sampling bias (Lin and Peddada 2020), further confirmed that bacterial differences between HAHT and natural sources were driven primarily by *Proteobacteria* (enriched in HAHT) and *Firmicutes* (enriched in natural) across all stages (Figure 3a,c, Figure S6- S12, p < 0.001). For fungi, differential abundance patterns were driven by *Ascomycota* species (Figure 3b,d, Figure S64-S12, p < 0.001). Notably, while baseline soil showed a large number of differentially abundant OTUs across a number of diverse phyla, the overall number of differentially abundant OTUs dramatically decreases through the IMO process, with the remaining differentially abundant phyla being primarily restricted to these clades (Figure 3a-d, Figure S6-S12, p < 0.001).

**Figure 3.**
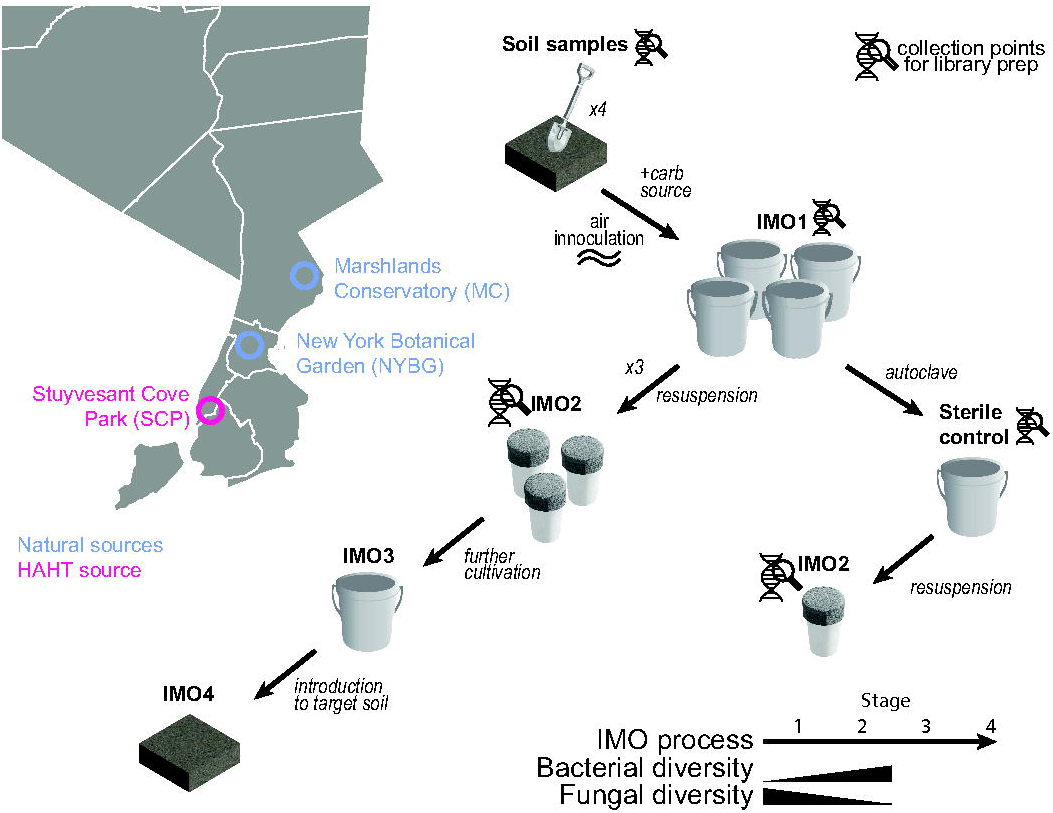
Differential abundance is driven by *Proteobacteria*, *Fermicutes*, and *Ascomycota*. ANCOM-BC results for differentially abundant microbes (p < 0.001) between HAHT and natural (a, b) soils (top 15 +/- differentially expressed OTUs) and (c, d) IMO2 day 7 cultures. Note, taxa are labeled using family or genus, while full OTU information may be found in Figure S4-S10. Relative abundance for (e) *Proteobacteria* and (f) *R. oryze* across IMO stage demonstrate how the IMO process enriches for certain OTUs. Alt text: Multi-panel graphic containing four horizontal bar charts (a through d) and two box plots (e and f). Charts a, b, c, and d show log fold change values spanning from negative to positive for specific microbial families and genera, color-coded by phylum legends in the center. Plots e and f show relative abundance trends on the y-axis from 0 to 1.00 across four timeline stages on the x-axis, categorized by location (MC, NYBG, SCP). Visually, the box plots show relative abundance sharply increasing after the baseline stage and remaining elevated through the final IMO2 day 7 stage.

We then further examined the dynamics of the most highly abundant clades: *Proteobacteria* for 16S libraries, and *Rhizopodacae* for ITS libraries. As observed previously, both HAHT (baseline vs IMO, p < 1e-6) and natural cultures (baseline vs IMO, p < 1e-8) show increased dominance by *Proteobacteria* (Figure 3e). Notably, the relative abundance of *Proteobacteria* increased dramatically regardless of sample origins. Similarly, *R. oryzae* appear to dominate over time, as highlighted by the Marshlands Conservancy samples. While marshland soil uniformly shows nearly 50% prevalence of *Ascomycota* (also known as sac fungi) with only a low prevalence of *R. oryzae*, these cultures are dominated by *R. oryzae* over time (baseline vs IMO, p < 1e-7, Figure 3f).

### Community structure revealed by PCoA

Two observations, 1) the decrease in number of differentially abundant OTUs, and 2) the dominance of differential abundance by a small number of phyla, suggests the primary effect of the IMO process is the cultivation of a limited number of microbes. However, like other fermentation products like wine, beer, and cheese, the diversity and presence of relatively low-abundance microbes may have large effects on the final product (Wolfe and Dutton 2015). Given that the IMO process is thought to preserve and propagate microbial diversity (Cho and Cho 2010, Reddy 2011), a process that can occur independently of the relatively large abundance of a few key microbes, we further examined microbial community structure using a Principal Coordinates Analysis (PCoA) using the Jaccard Index. Given the relatively high abundance of a few species in our previous results, we chose to utilize the Jaccard Index as it is an unweighted measure of community similarity that is robust to differences in abundance. The resulting PCoA figures demonstrate a striking difference between bacterial and fungal community structures of soil samples of different origins (Figure 4a,b). Samples obtained from the HAHT Stuyvesant Cove Park cluster tightly and distantly from the cluster of samples obtained from natural settings. Further, all sterile samples comprise their own cluster. This relatively strong clustering based on sample origin suggests that the IMO process manages to preserve some degree of community similarity.

**Figure 4.**
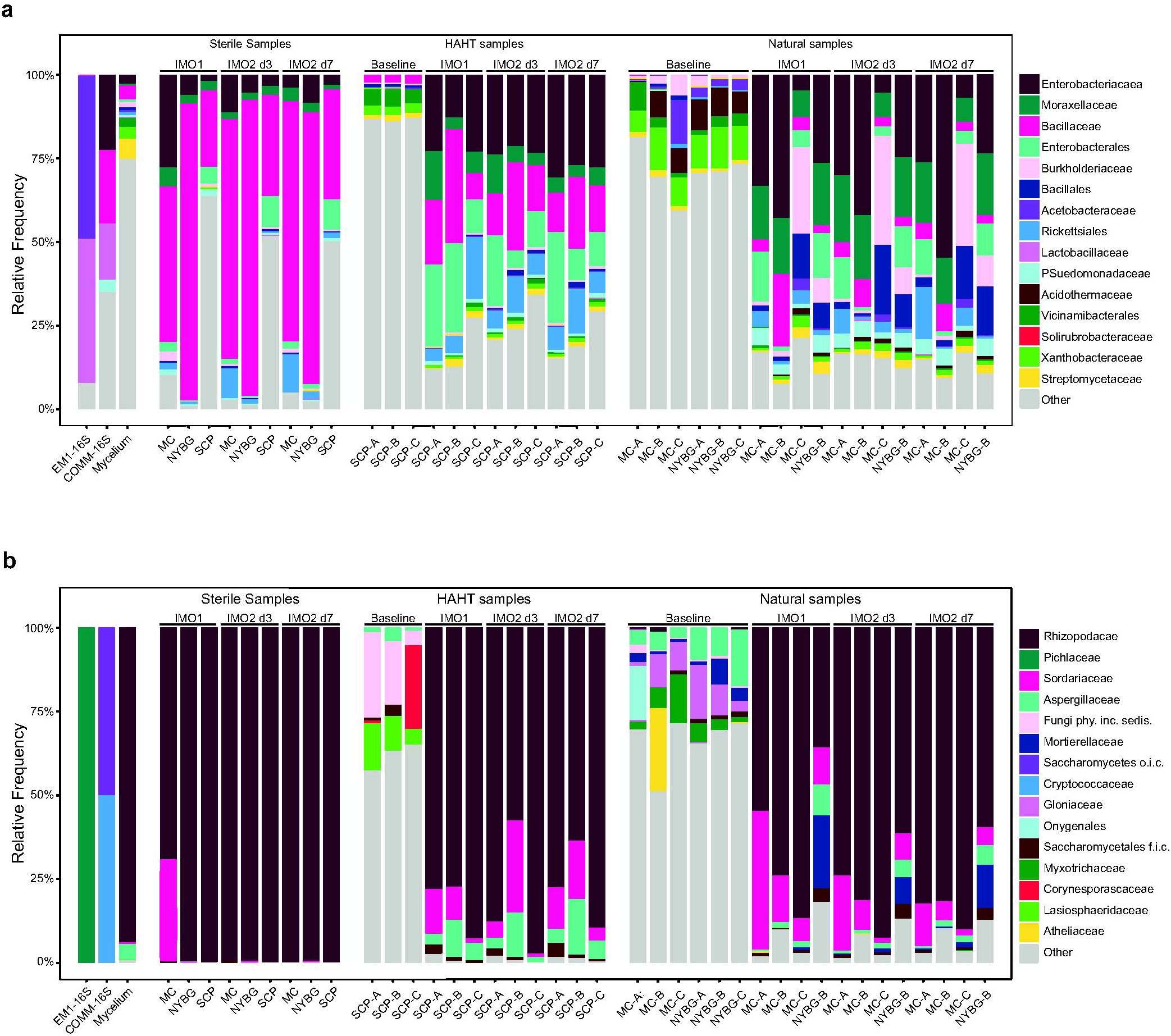
Community structure and diversity during the IMO process. (a-d) 16S and ITS community structures reflect initial soil microbiomes and IMO stage in PCoA visualizations (Jaccard Index). (e, f) β-diversity (Unweighted UniFrac) of bacterial communities initially increases over the IMO process while β-diversity of fungal communities decreases over time Alt text: Multi-panel figure containing four three-dimensional principal coordinate analysis (PCoA) scatter plots (panels a through d) and two stacked box plot graphs (panels e and f). The PcoA scatter plots on the left show colored data points clustering into distinct groups based on soil type (natural, HAHT, sterile, control) and experimental stage. The right-hand box plots measure Unweighted UniFrac Distance on the y-axis across four timepoints on the x-axis, from baseline to IMO2 day 7. Visually, the box plots show beta-diversity values trending slightly upward for the 16S bacterial communities in panel e, while trending downward over time for the ITS fungal communities in panel f.

Examination of the 16S PCoA results show four distinct clusters: natural, HAHT (i.e. sourced from Stuyvesant Cove Park), sterile-sourced, and synthetic (i.e. EM1 and Zymo Community Standards). Notably, our single primarily- mycelium sample (Methods and Materials) showed a remarkable degree of bacterial community structure that clustered closely with HAHT samples, reflecting the fact that the mycelium sample was pulled from the HAHT IMO1 box. Interestingly, Principal Coordinate 3 (∼5% of variance) shows a strong correlation with IMO stage in the 16S data, but not Principal Coordinates 1 and 2 (Figure 4c).

Examination of the ITS PCoA results show a lower level of stratification between inoculant sources. While all four sources may be distinguished, the overall distance between ITS community profiles is attenuated. Unlike in the case of the 16S PCoA results, a strong stage-specific effect is noticeable on the first two principal coordinates of the ITS data (Figure 4d). In the case of both HAHT and natural data sets, soil samples visibly cluster together while the microbial communities of later IMO stages appear increasingly distant from the initial soil inoculant samples. The ITS PCoA results also demonstrate how the forest-inoculated samples cluster closely with marshland samples but remain very distant from the sterile, HAHT, and synthetic communities (Figure 4b). This absence of strong distinguishing factors between the singular forest sample and our biologically triplicate marshland samples thus supported our choice of including forest and marshland samples in a single classification of natural samples for all downstream analyses.

Interestingly, both 16S and ITS PCoA figures demonstrate a tripartite structure in the first two principal coordinates where sterilized controls appear in somewhat intermediate positions between natural and HAHT sources (Figure 4a,b). This tripartite structure remains true regardless of IMO stating, thus demonstrating how inoculation source plays a larger role in defining microbial composition than growth media. The absence of strong stage-specific effects along the first two 16S principal coordinates thus suggests that the first two stages of the IMO process, in a manner similar to standard LB growth media, strongly determine bacterial community composition. However, this same conclusion does not hold for fungal communities, as stage-specific effects are readily apparent in the first two principal coordinates of the ITS analysis. This difference in stage specificity is consistent with prior observations demonstrating how fungal community succession produces more deterministic responses to external factors, while bacterial community succession patterns appear to be more random.

### Diversity dynamics during cultivation

While the previous results demonstrate how the early stages of the IMO process are dominated by a limited number of taxa, it is still possible that the early stages of the IMO process can retain high levels of diversity for later transfer. To further test whether the initial steps of the IMO process can maintain microbiome diversity prior to eventual microbiome transfer, we calculated the β-diversity (Unweighted UniFrac) both within and between natural and HAHT samples at the soil, IMO1, and IMO2 (day 3 and day 7) stages. Similar to the Jaccard Index, the Unweighted UniFrac measure is robust to changes in relative abundance. We found that sample source (i.e. HAHT vs natural vs sterile) were significant factors in determining unweighted UniFrac values (PERMANOVA, p < 0.05) when compared in aggregate across all stages and within each stage for both 16S and ITS libraries (Figure 4e,f). We then examined the distribution of pair-wise UniFrac values for a more detailed analysis of the interaction between stage and source.

As expected, the within-group bacterial (PERMANOVA, p < 0.001) and fungal (PERMANOVA, p < 0.001) UniFrac estimates for natural soils were higher in natural soils than in HAHT soils (Figure 4e,f). While the inclusion of the single forest sample with the three marshlands samples likely increased the distribution of pair-wise UniFrac computations, both the minimum and lower quartile values still remain well above the UniFrac distribution for HAHT soils. The magnitude of this difference, which persists over all IMO stages tested in our study, suggests that the conclusions drawn from our within-group comparisons are robust to the inclusion of the forest sample.

Interestingly, the early stages of the IMO process appear to increase within-group 16S diversity (PERMANOVA, p < 0.05) but decrease within-group ITS diversity (PERMANOVA, p < 0.001)(Figure 4e,f). Specifically, the 16S diversity between samples during the IMO1 and IMO2 stages is higher than in soil. This holds true for within-group samples with a natural origin as well as for HAHT origins. There is a noticeable difference in within-group 16S diversity when moving from soil to the IMO1 stage. However, the IMO1 stage, which is cultured on rice, and the IMO2 stage, which is cultured on unrefined jaggery sugar, do not appear to have noticeably different levels of 16S diversity regardless of origin. The overall effect is that once the soil microbiome is transferred to a carbohydrate medium, the resulting bacterial diversity is maintained over the time period that we employed in this study. This suggests that after an initial period of selection, the resulting bacterial communities appear to be relatively stable. Alternatively, the within-group ITS diversity continuously trends downward from the soil stage to the IMO2 stage, both for natural and HAHT samples. This suggests that the IMO process is not an effective method for propagating fungal communities.

Surprisingly, the stability of within-group HAHT and natural 16S diversity contrasts with sterilized media inoculant (e.g. potentially airborne inoculated), where both bacterial (PERMANOVA, p < 0.001) and fungal (PERMANOVA, p < 0.001) within-group diversity appears to decrease by the IMO2 day 7 stage. The initial within-group 16S diversity for sterile IMO1 and IMO2 day 3 lies between the 16S diversity of the natural and HAHT samples but decreases below the 16S diversity levels present in the HAHT sample by the IMO2 day 7 stage. Alternatively, the within-group ITS diversity for the sterilized media remains at a very low level throughout the IMO process. This observation potentially results from a resource-limited carrying capacity of the sterilized media IMO2 culture, though it is unclear why such a carrying capacity may be reached earlier in these cultures vs HAHT and natural IMO2 cultures. This difference in carbohydrate metabolism is likely related to differential abundances of *Proteobacteria* and *Firmicutes* spp., as *Proteobacteria* generally metabolize simpler sugars at a higher rate than *Firmicutes* (Bradley and Pollard 2017, Méndez-Salazar *et al*. 2018). Analysis of predicted metagenome functions using MetaCyc annotations in PICRUSt2 (Table S4) shows a significant enrichment of carbohydrate metabolism in IMO cultures in comparison to soil samples (Figure S13). For example, the list of 336 pathways that were differentially abundant in both HAHT and natural cultures (adjusted p-value < 0.05, Table S5) include the superpathway of glucose and xylose degradation (PWY-6901), glucose degradation (oxidative) (DHGLUCONATE- PYR-CAT-PWY), glucose and glucose-1-phosphate degradation (GLUCOSE1PMETAB-PWY), lactose and galactose degradation I (LACTOSECAT-PWY), sucrose degradation III (sucrose invertase)(PWY-621) and IV (sucrose phosphorlyase)(PWY-5384). As these results are based on predictions, further validation of hese metabolomic changes require a different shotgun metagenomic sequencing strategy. Interestingly, such resource limitation and altered community metabolomes may also suggest the possibility of further complex, successional dynamics beyond the relatively short time window studied here (Wolfe *et al*. 2014, Wolfe and Dutton 2015).

As expected, 16S and ITS soil diversity appears to be highest when comparing between HAHT and natural groups. Between-group ITS diversity appears to decrease over the early stages of the IMO process, suggesting the presence of environmental factors that are gradually homogenizing fungal communities by favoring the growth of certain microbes. While such loss of between-group ITS diversity is expected as a result of decreasing within-group ITS diversity, between-group 16S diversity levels are surprisingly constant through the IMO process. This observation highlights the seemingly paradoxical observation that the IMO process results in an increase of the within-group 16S diversity.producing greater species/taxonomic diversity than what is present in the inoculant. Alternative possibilities must be considered in order to explain the later detection of bacterial species not originally detected in soil samples.

One possibility is that this increase is resulting from sampling error resulting from biological or technical limitations (see Methods and Materials). This explanation is unlikely given the extremely high read depths produced by AVITI sequencing – UniFrac computations were performed at a 280000 read sampling depth, with many libraries producing over 10x that number of reads (Table S3). Rarefaction analyses and plots for unweighted UniFrac values show Spearman correlation coefficients all in excess of ρ=0.99 for both 16S and ITS data sets at this sampling depth. Instead, the increase in bacterial diversity during IMO cultivation may have potentially resulted from airborne inoculation, consistent with intermediate within-group 16S diversity levels in sterile controls. This stands in contrast to the within-group ITS diversity for the same controls, which begins and remains at low levels. Rapid recolonization by species that escaped sterilization also remains unlikely, as the resulting diversity should be a subset of the diversity already existing in our live, unsterilized soil samples; this does not appear to be the case.

## Discussion

Recent work has sparked increased interest in understanding the relationship between soil amendment, soil health, and the underlying microbiome structures of amended soil, particularly as it relates to crop yield and land management (reviewed in (Hao and Ashley 2021, Hermans *et al*. 2023)). Results from prior studies in understanding the relationship between soil amendment and shifts in microbiome community structure remain mixed, particularly given the intrinsic coupling of environmental conditions (e.g. nutrient availability and physiochemical properties of soil) and microbial growth (Liu S *et al*. 2021, Liu W *et al*. 2023, Oberholzer *et al*. 2024). For example, one study amended soil using compost treated with either the commercially available “effective microorganism” EM1 amendment or a single-species *B. subtilis* amendment, finding that both treatments increased bacterial diversity but had no effect on fungal diversity. Importantly, these changes were found to been mostly likely driven by soil physiochemical properties (Liu S *et al*. 2021). This result is supported by a meta-analysis by (Liu W *et al*. 2023), where alterations of microbial biomass resulting from organic soil amendment were likely driven primarily by nutrient availability and carbon content. A further study attempting to directly separate EM1 amendment-driven and substrate-driven effects on soil microbiomes found no consistent effect of EM1 treatment on the physiochemical properties of the treated soil, including soil pH, nutrient availability, and more. Direct alteration of soil microbiome composition from EM1 treatment was also found to be detectable but weak only under treatment conditions far exceeding the commercially recommended treatment levels (i.e. 100x recommended dose). Importantly, this 100x dosage also induced significant alterations of the underlying soil substrate, altering the physiochemical properties of the treated soil (Oberholzer *et al*. 2024).

Similarly to the commercially available EM1 treatment, the overall goal of the IMO process is to improve the health of soil through the transfer and/or downstream activity of certain microbial communities. In parallel with previous work directly testing the effects of soil amendment on microbial community structures, our study provides the first comprehensive characterization of microbial community dynamics during Korean Natural Farming practices across multiple ecological contexts. In particular, our results demonstrate that the early stages of IMO cultivation are primarily dominated by a small number of well-characterized microbes. Despite this, we found that distinct patterns emerge based on initial inoculum source and sterilization treatment. Analyses that are robust to differences in abundance, i.e. the Jaccard Index and the Unweighted UniFrac, yield distinct, striking results. We observed that bacterial communities in HAHT environments contain lower microbial diversity than natural communities, both at baseline and following IMO treatment, regardless of original inoculation source. This result highlights the importance of preserving historic soils as a source of biodiversity, as recently HAHT soil environments do not easily replicate their natural longstanding counterparts.

One potential benefit of IMO treatment would be in utilizing the inoculation of soils in HAHT environments in order to diversify resident microbial communities. However, our limited results from the early stages of the IMO process suggest that it is unlikely that the IMO process could help immediately increase fungal diversity, as the apparent fungal diversity decreased over successive cultivation steps. It is important to note that our results are subject to known biases inherent to amplicon-based sequencing, including preferential amplification resulting from variable rRNA copy number or primer bias (reviewed in (Bálint *et al*. 2016)). We also note that apparent reductions in taxonomic diversity may not necessarily correlate with functional diversity (Graco-Roza *et al*. 2021, Li *et al*. 2021), highlighting the need for follow-up studies measuring underlying both the functional and taxonomic diversity of fungal communities during IMO cultivation.

Additionally, our results suggest that the IMO process generally selects for specific taxa adapted to the cultivation conditions, though some degree of bacterial diversity can be maintained over time. It is important to note, however, that the microbial community dynamics characterized in this study remain restricted to early-stage IMO cultivation, and as such, the later stages of IMO cultivation could still significantly affect microbial diversity, ecological function, and perhaps most importantly, plant-microbe interactions. Regardless, the persistent differences between HAHT and natural microbial communities throughout the early stages of IMO cultivation do suggest that community succession may potentially require longer timeframes than those examined in this study, or that initial community composition strongly constrains microbial community dynamics. This finding has practical implications for soil bioremediation efforts, indicating that inoculum source remains important even after multiple cultivation steps. It remains to be seen how microbial communities shift through the “IMO3” and “IMO4” stages. There remains the possibility that, rather than cultivating a large consortium of fungal species, the IMO process propagates a few key players that then create ideal conditions for future biodiversity.

The observation that bacterial diversity was maintained or increased during cultivation suggests that early-stage IMO fermentation may facilitate airborne inoculation in addition to soil-derived propagation. While this may broaden bacterial diversity, it also challenges the notion that IMOs exclusively preserve “indigenous” microbial communities specifically from the uppermost layers of the soil horizon (Cho and Cho 2010, Reddy 2011).

Taken together, these findings suggest that the early stages of IMO cultivation functions less as a direct transfer of indigenous soil microbiota and more as a selective enrichment system shaped by cultivation conditions and environmental inoculation. While it is possible that the claimed benefits of the IMO process and the increasing popularity of KNF practices results from the introduction of raw nutrients or keystone species for further succession, not necessarily from overall microbiome transfer, downstream studies testing longer cultivation times, soil reinoculation outcomes, and plant health effects will need to be performed to confirm this hypothesis. Further longitudinal studies should be performed to characterize the microbial community dynamics in the later stages of the IMO process – most importantly after reintroduction of IMO products into soil. Importantly, understanding the relative contribution of the physiochemical properties of soil substrate vs. microbial community structure will be critical for a full understanding of the IMO process. This study also presents the hypothesis that airborne inoculation of IMO cultures may be an important, but underappreciated factor in determining microbial composition during the IMO process; further air sampling controls would be critical in confirming this hypothesis. A fuller functional metabolomic e.g. shotgun metagenomic or metatranscriptomic, characterization of how microbial community structure utilizes available nutrients to alter its local environment should also contribute key insight into our fundamental understanding of how long-term ecological stability is achieved in soils.

## Acknowledgements

The authors would like to thank Michael Gambino, Kristen Pareti and Leah Cass from Marshland’s Conservancy, Eric Sanderson, Elliot Nagele and John Zeiger from New York Botanical Garden, Sara Evans from Greenwood Cemetery, Dina Elkan at Solar One, Biotech Without Borders, the Rockefeller University Genomics Resource Center (RRID: SCR_020986), and thought partners/community collaborators Brooke Singer, Sneha Ganguly, Craig Trester, Jason Sinopoli, Blacki Migliozzi, Nathan Hunter, Journei Bimwala, Rodney Santiago, Andrea Estrada, Margaret Boozer, Andie Marsh and Sara Perl Egendorf for their contributions to this project. The authors would like to thank Landscape Architecture Design Team of Mathews Nielsen Landscape Architects, P.C. for providing soil specification data. The authors would like to thank the Urban Soils Institute and George Lozefski, Igor Bronz, Margaret Boozer, Paul Mankiewicz, Rich Shaw, and Maha Deeb for their helpful discussions regarding soil composition.

## Data Availability

Raw sequencing data is deposited in the NCBI Sequence Read Archive as PRJNA1456652. Analysis scripts, metadata, and QIIME2 viewer files for PCoA and taxa bar plots are available at https://github.com/ulee-sciscripts/imo.

## Author Contribution Statement

CT contributed to: conceptualization, funding acquisition, investigation, methodology, writing original draft and reviewing and editing. SM contributed to: formal analysis, investigation, methodology, visualization, and writing original draft and reviewing and editing. EP contributed to: investigation and methodology. UL contributed to: conceptualization, data curation, formal analysis, funding acquisition, investigation, methodology, project administration, resources, software, supervision, writing original draft and reviewing and editing.

## List of Supporting Materials Legends

**Table S1 Observations during collection process.** Data recorded during IMO1 installation, IMO1 retrieval, and IMO2 processing.

**Table S2 Physical qualities of soil samples.** Results of commercial soil analysis from single MC, NYBG, and SCP soil samples

**Table S3 Summary metrics for AVITI sequencing.** Index assignment and metrics provided for sequencing.

**Table S4 Functional pathway predictions for 16S libraries.** Results of PICRUSt2 functional pathway predictions for IMO libraries.

**Table S5 Significant differentially abundant pathway predictions.** Pathways that are significantly differentially abundant between stages in both HAHT and natural IMO cultures, adjusted p-value < 0.05.

**Figure S1 Experimental process of IMO cultivation.** Representative photographs show the IMO cultivation process used in this study for (a-c) live culturing of IMO from untreated soil and (d-e) sterilized soil. (f) Resulting IMO1 cultures were used to generate IMO2 cultures

Alt text: Six photographs labeled a through f showing physical steps of Indigenous Microorganism (IMO) cultivation. Panel a shows a mesh-covered wooden box filled with white rice placed on open soil. Panel b shows a similar box buried under leaves and a thin white mesh fabric outdoors. Panel c shows a box enclosed within wire mesh fencing. Panel d displays two plastic storage bins with small ventilation holes stacked on a wooden shelf indoors. Panel e features a close-up of a gloved hand holding a block of dense, white fungal mycelium growing on rice grain substrate. Panel f shows glass jars containing dark organic mixture covered with paper towels on a laboratory workspace.

**Figure S2 Alpha diversity metrics for IMO libraries.** (a, b) 16S and (c, d) ITS alpha diversity metrics by (a, c) stage and by (b, d) source.

Alt text: Four box plot graphs arranged in a two-by-two grid displaying alpha diversity metrics (Faith PD) on the y-axis. The top two charts measure 16S bacterial alpha diversity, and the bottom two charts measure ITS fungal alpha diversity. The left-hand plots show diversity metrics changing across five stages on the x-axis: Soil, Baseline, IMO1, IMO2 day 3, and IMO2 day 7, showing an immediate drop from soil to baseline before stabilizing. The right-hand plots subdivide the data by geographic sampling sources along the x-axis: Control, Marshland Conservancy, New York Botanic Garden, and Stuyvesant Cove Park, showing variable box spreads and individual outlier points.

**Figure S3 PCoA by source.** PCoA for (a, b) 16S and (c, d) ITS libraries colored by source.

Alt text: Four three-dimensional principal coordinate analysis (PCoA) scatter plots arranged in a two-by-two grid labeled a through d. The top two plots (a and b) show 16S bacterial library data, while the bottom two plots (c and d) show ITS fungal library data. Each plot features colored circular data points distributed across three axes. A single shared legend at the bottom maps the data point colors to their respective soil source types: grey for Control, blue for Natural, pink for HAHT, and yellow for Sterile. The points are visibly grouped into distinct, isolated clusters corresponding to these color designations across all four panels.

**Figure S4 16S diversity metrics by site origin.** UniFrac distances between 16S libraries for NYBG and MC samples.

Alt text: Four box plot charts labeled a through d arranged in a two-by-two grid. The charts plot distance metrics on the y-axis from 0 to 1.0 against various location sample groups on the x-axis, such as MC, NYBG, and SCP sub- sites and sterile controls. Each panel calculates distances relative to a specific reference baseline site, noted in the individual plot titles: panel a references MC-b, panel b references MC-a, panel c references MC-c, and panel d references New York Botanic Garden-b. The box plots across all panels are color-coded by location type, showing fluctuating median distance scores across the different comparative groupings.

**Figure S5 ITS diversity metrics by site origin.** UniFrac distances between ITS libraries for NYBG and MC samples.

Alt text: Four box plot charts labeled a through d arranged in a two-by-two grid format. Each chart displays distance values on a y-axis from 0 to 1.0 against comparative site sample groupings on the x-axis, color-coded by location type. The individual panels show 16S distance metrics relative to a specific reference baseline site noted in their top titles: panel a references New York Botanic Garden (b), panel b references Marshland Conservancy (b), panel c references Marshland Conservancy (c), and panel d references Marshland Conservancy (a).

**Figure S6-S12 Differential abundance (ANCOM-BC) between IMO samples derived from HAHT and natural source soils.** Full differential abundance results, p < 0.001. Plots are provided on separate pages in the following order: (5) 16S-soil, (6) 16S-IMO1, (7) 16S-IMO2d3, (8) 16S-IMO2d7, (9) ITS-soil, (10) ITS-IMO2d3, (11) ITS-IMO2d7. Note, ITS-IMO1 showed no significantly differentially abundant taxa.

Alt text: A horizontal divergent bar chart showing differential abundance. The x-axis measures Log Fold Change (LFC) fagainst a vertical list of specific bacterial Feature IDs on the y-axis. Bars extending to the right are colored blue to represent enriched taxa, while bars extending to the left are colored orange to represent depleted taxa.

**Figure S13 Significant differentially abundant pathway predictions.** Top 20 most significant differentially abundant pathways in (a) natural and (b) HAHT IMO cultures, adjusted p-value < 0.05.

Alt text: Two multi-chart panels labeled a and b displaying pathway predictions. Panel a is titled “Natural” and panel b is titled “HAHT”. Each panel consists of three side-by-side components: a vertical list of 20 metabolic pathways on the left, a multi-colored horizontal bar chart in the middle plotting Relative Abundance across stages (soil, imo1, imo2_d3, imo2_d7), and a solid blue horizontal bar chart on the right showing log2 fold change values ranging from 0.0 to 7.5 next to a vertical list of adjusted p-values.

