## Supplementary figures and images for "Korean Natural Farming practices are dominated by a limited number of microbes and decrease fungal diversity"

### FigS1

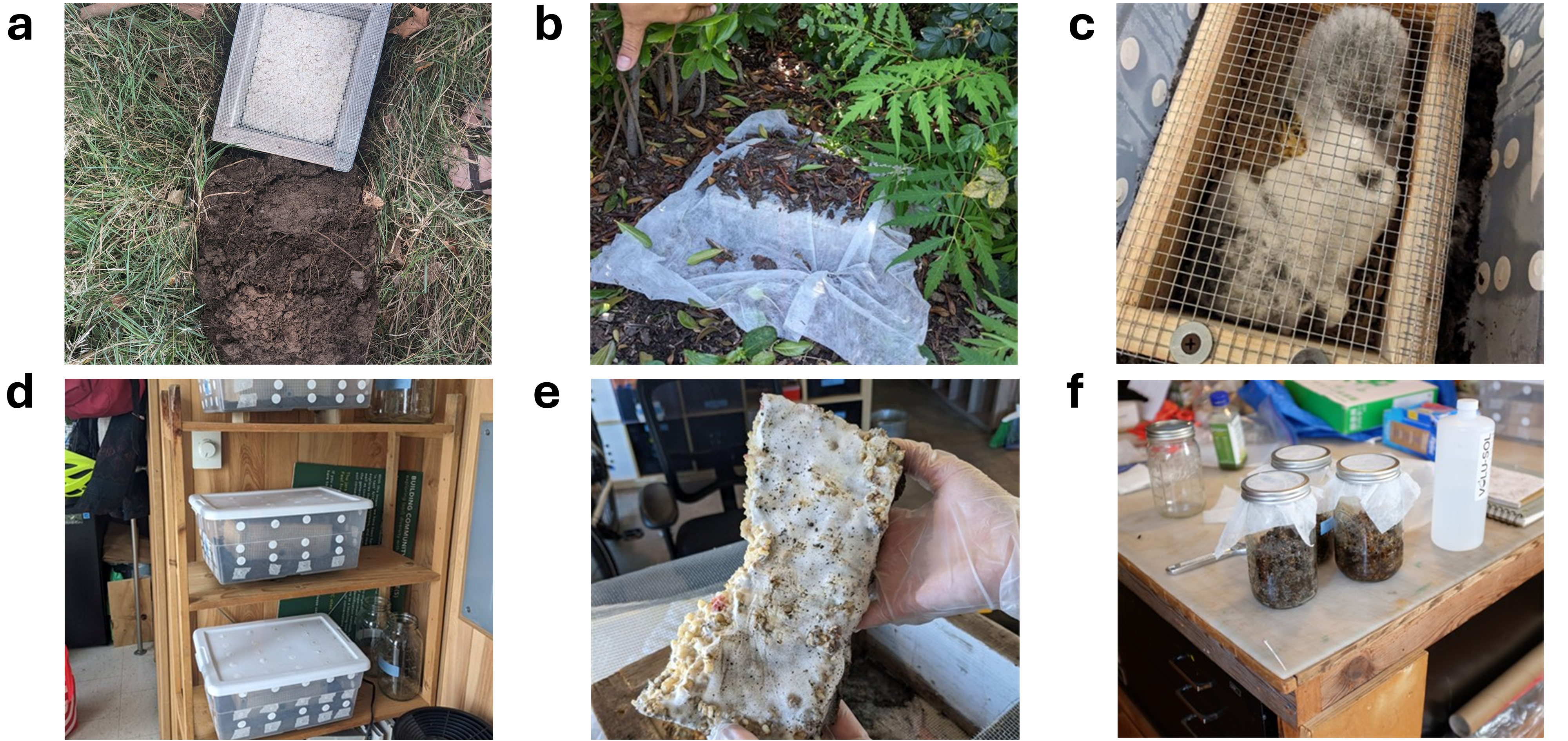

### FigS2

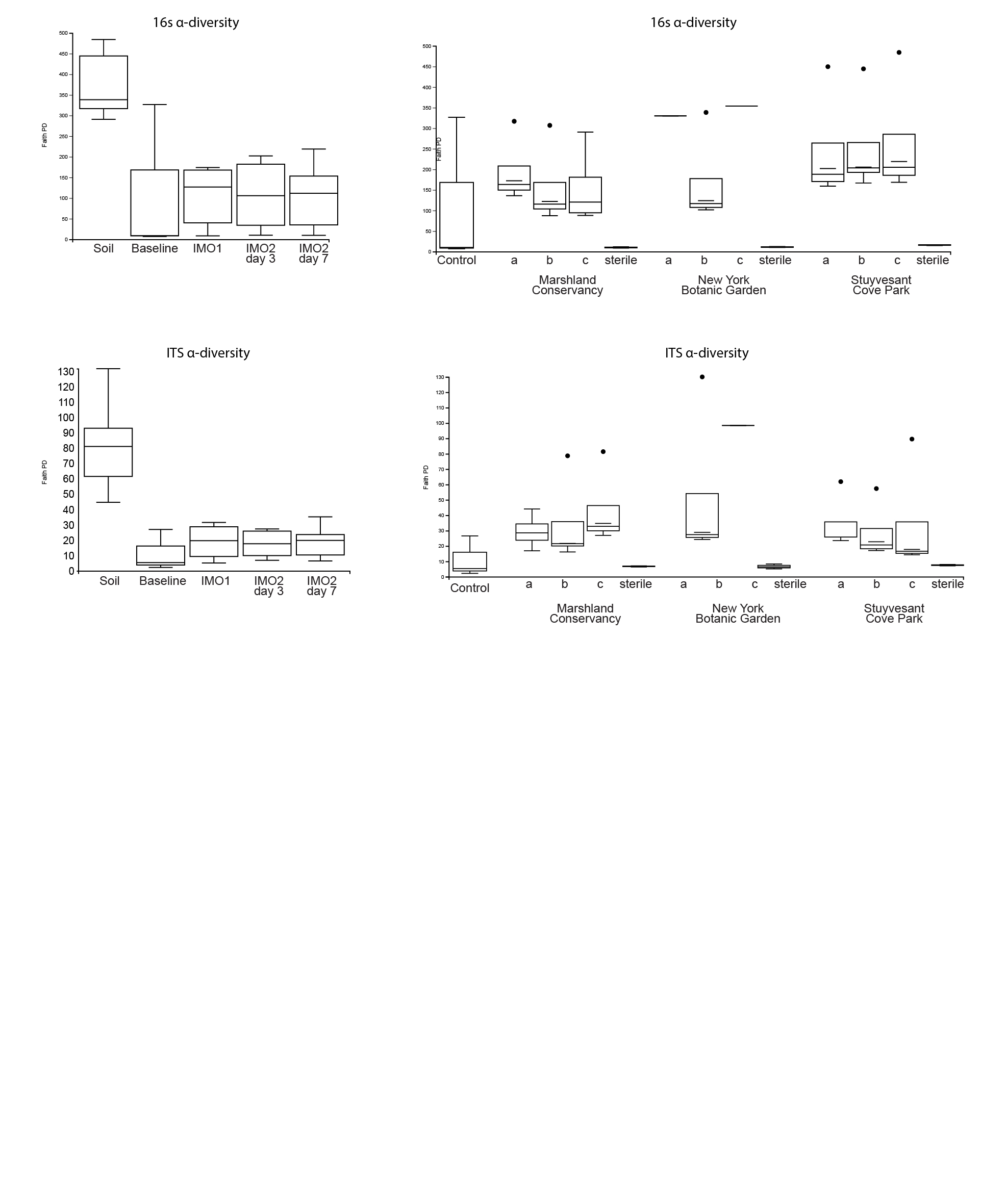

### FigS3

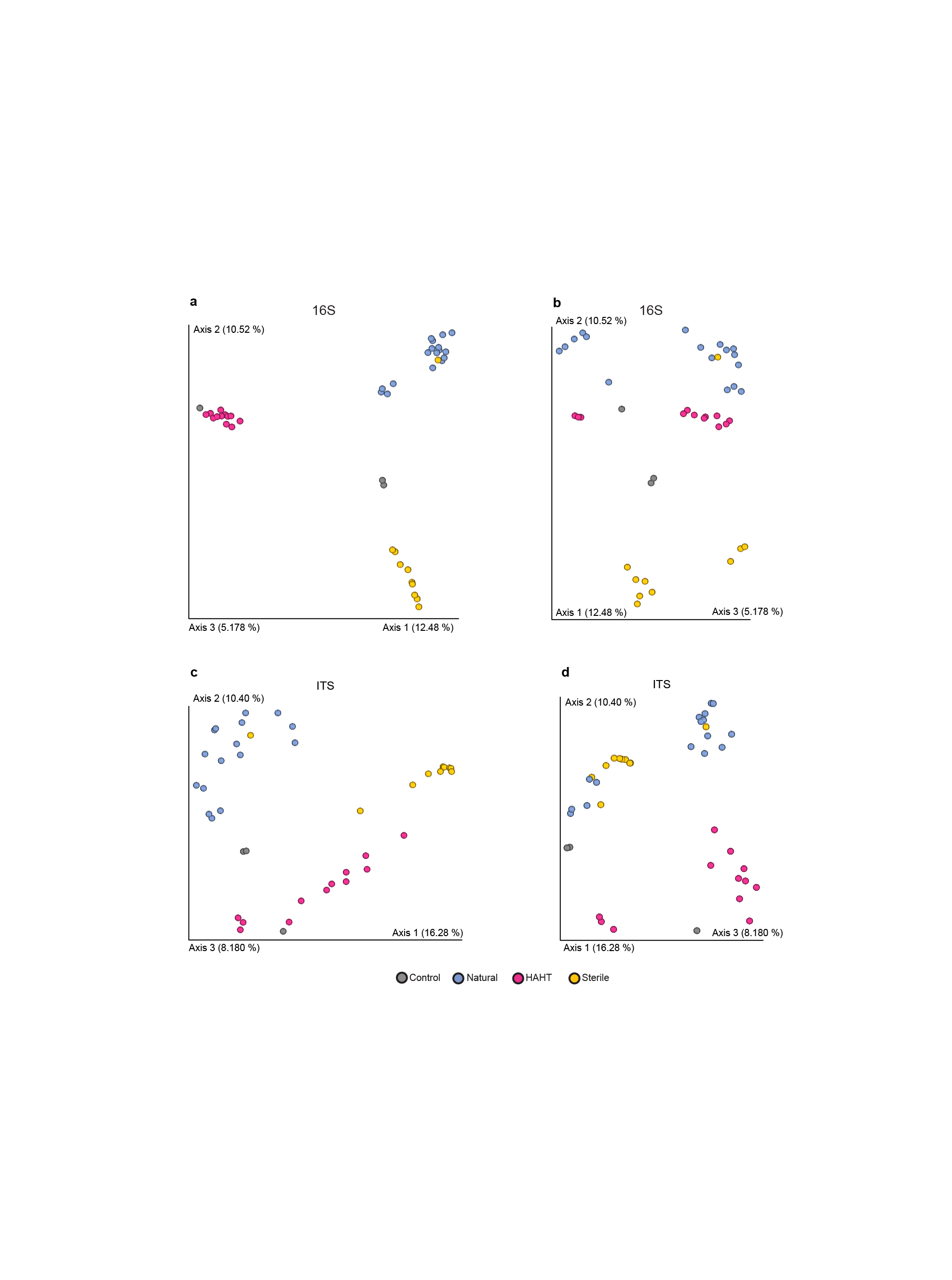

### FigS4

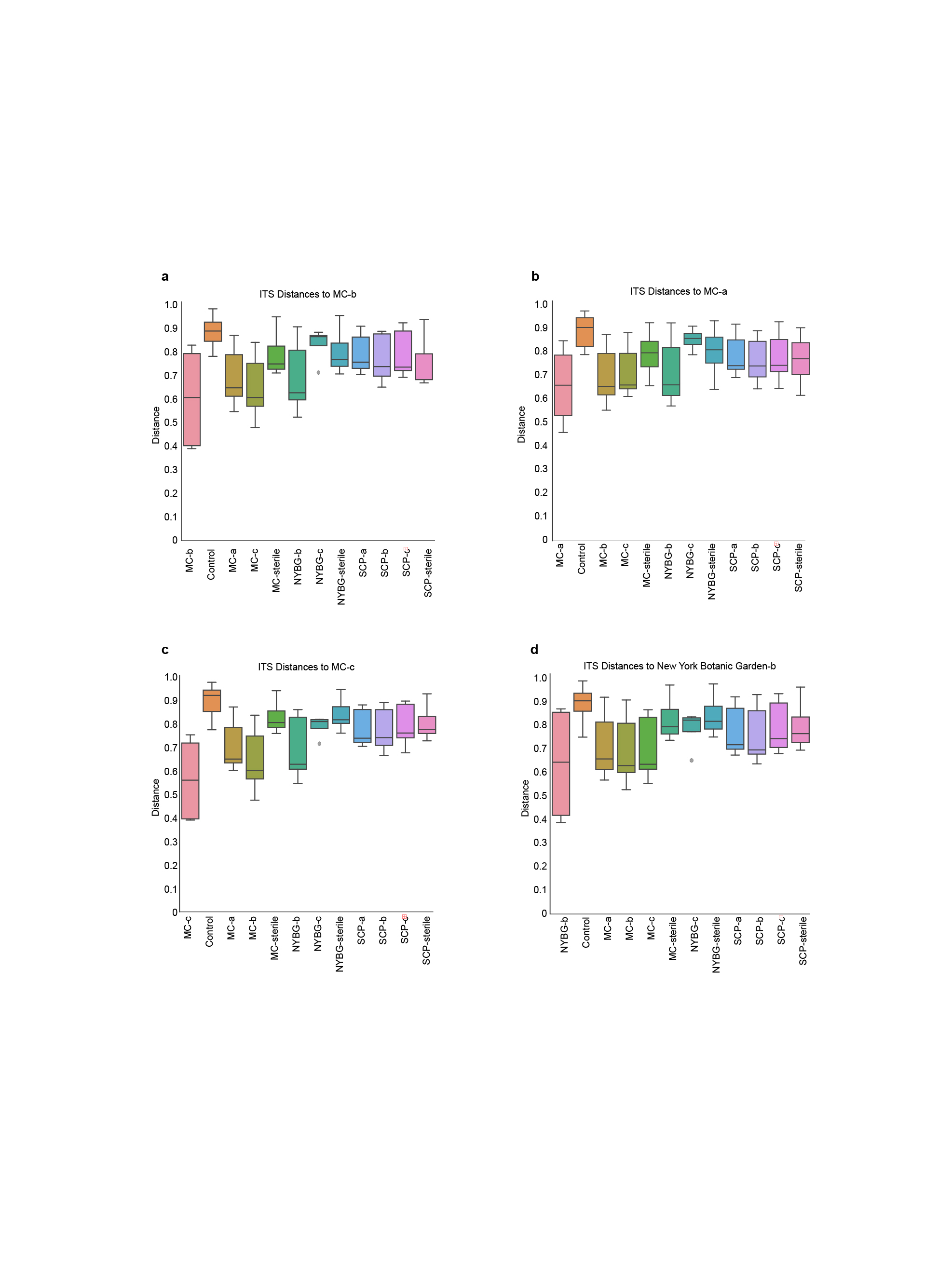

### FigS5

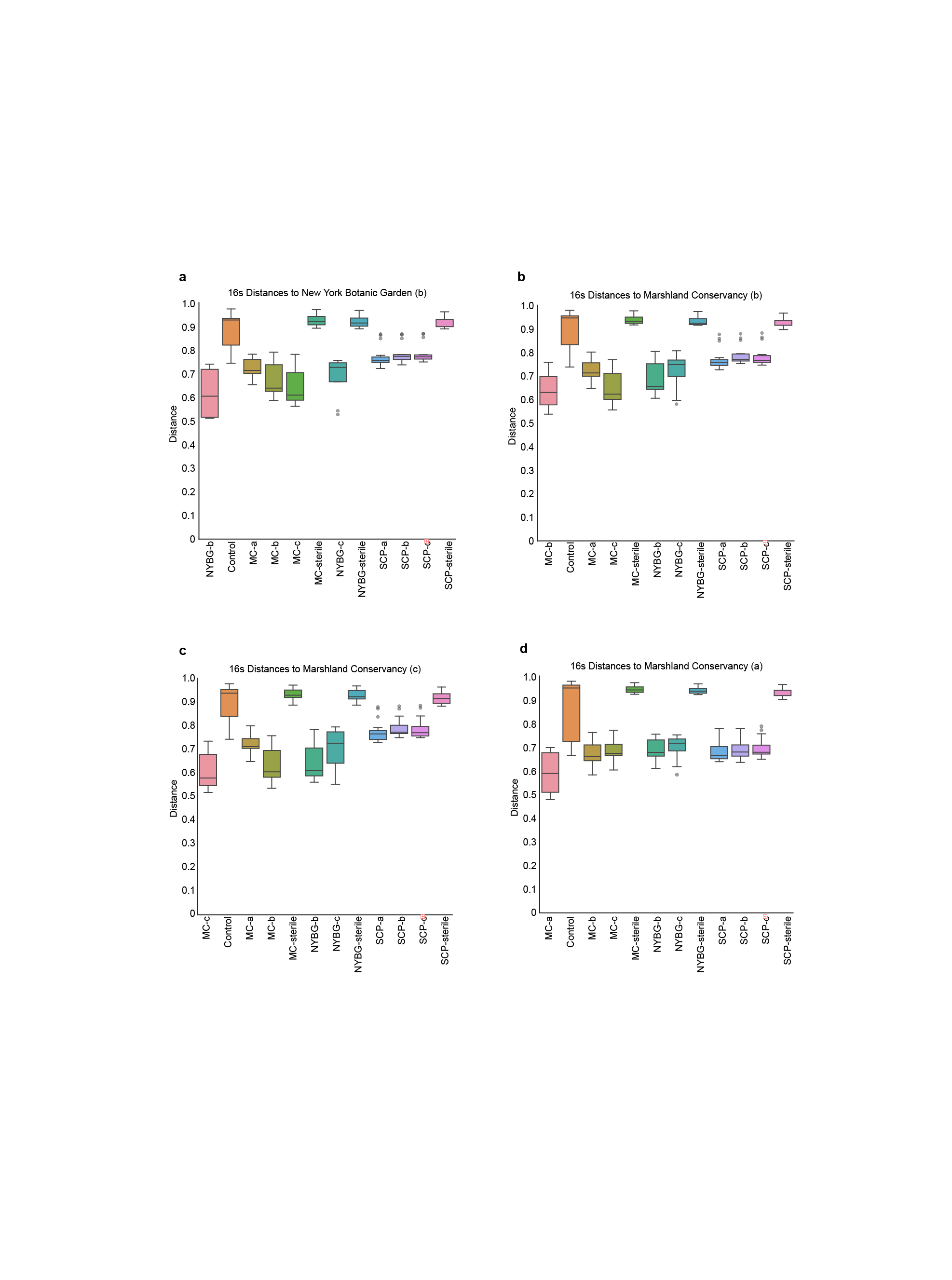

### FigS6

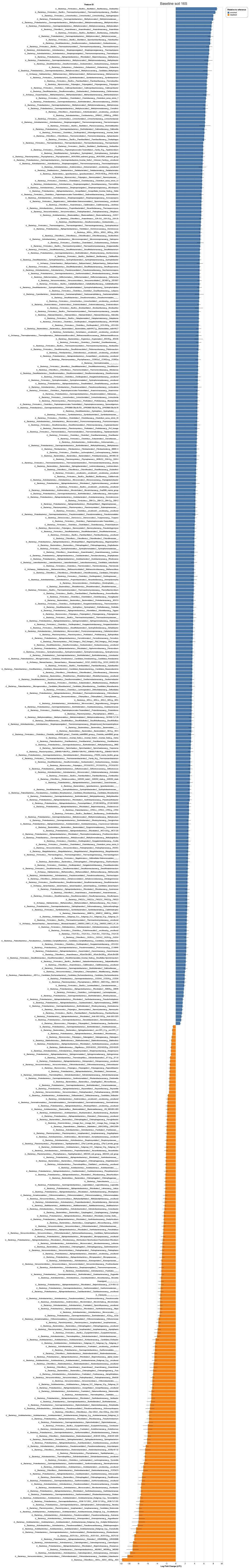

### FigS7

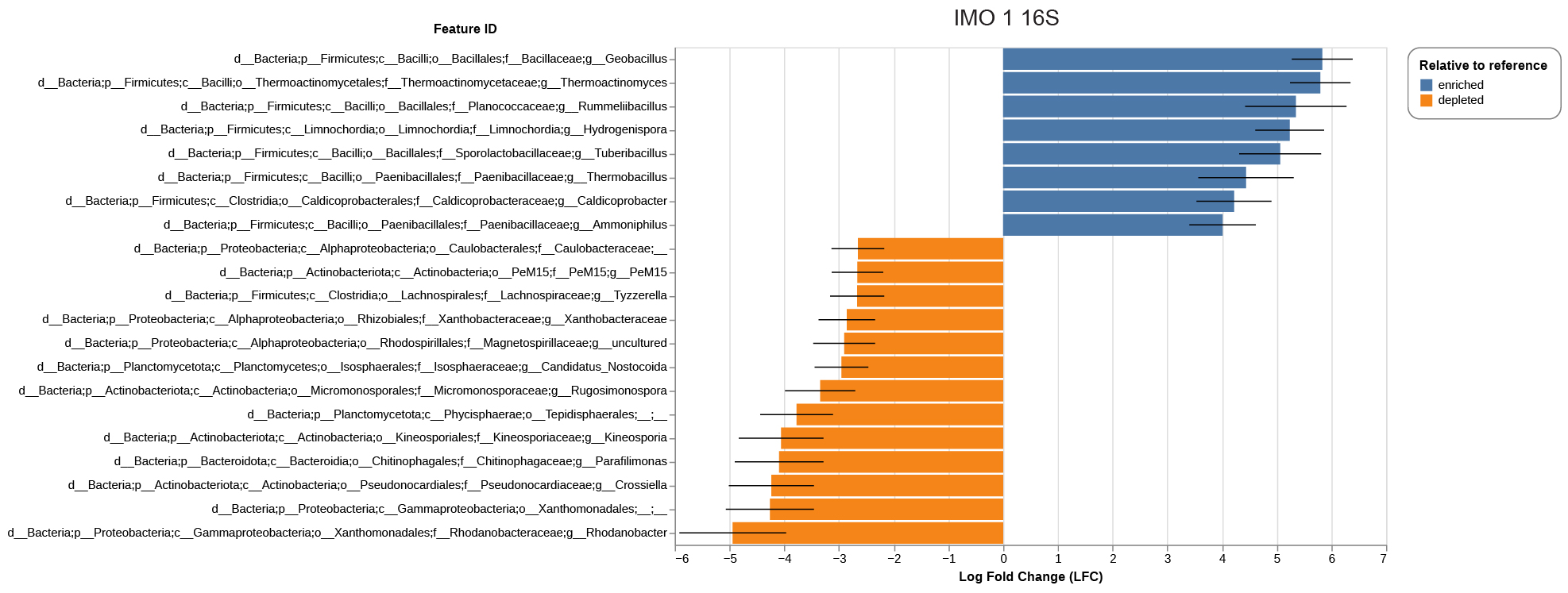

### FigS8

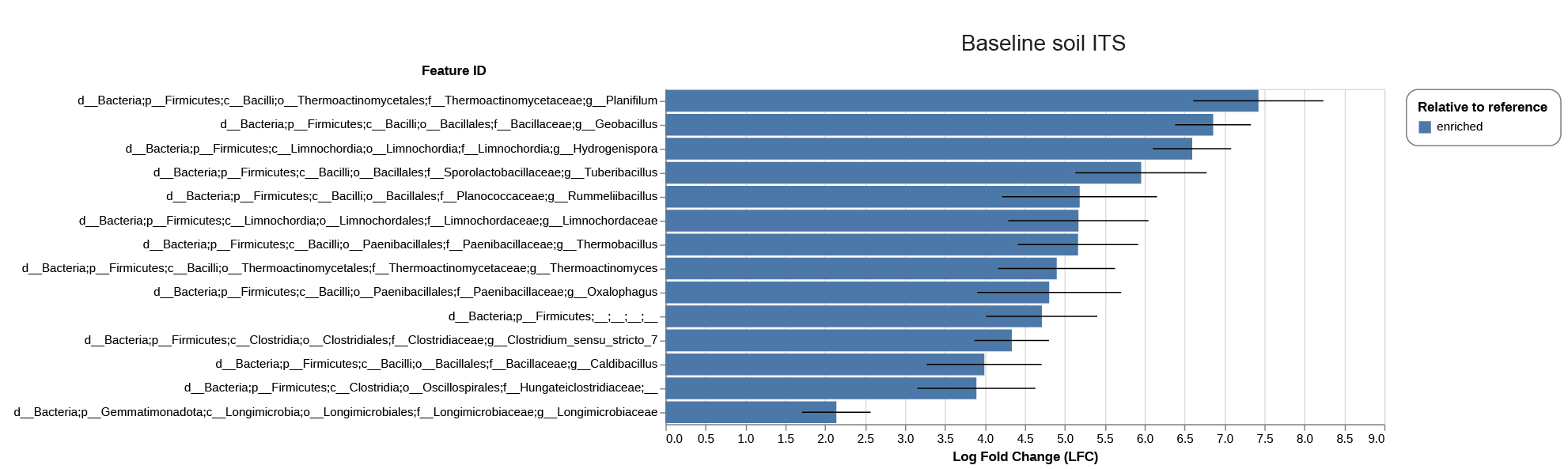

### FigS9

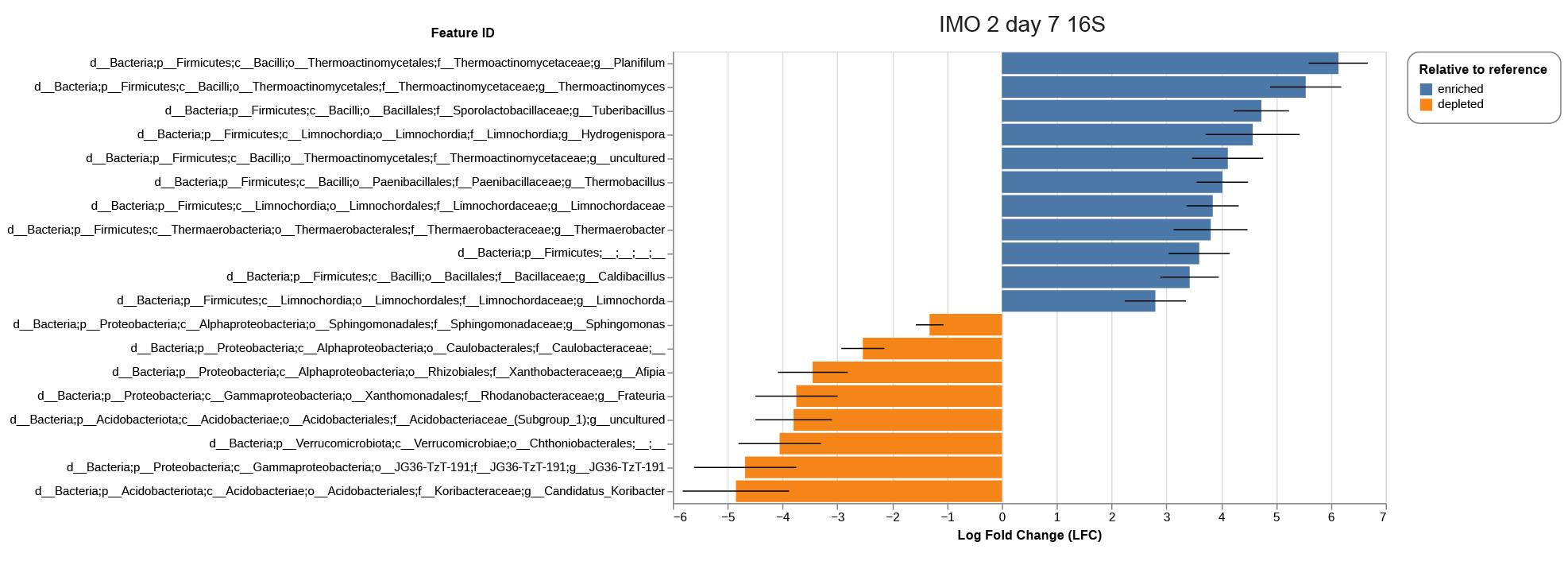

### FigS10

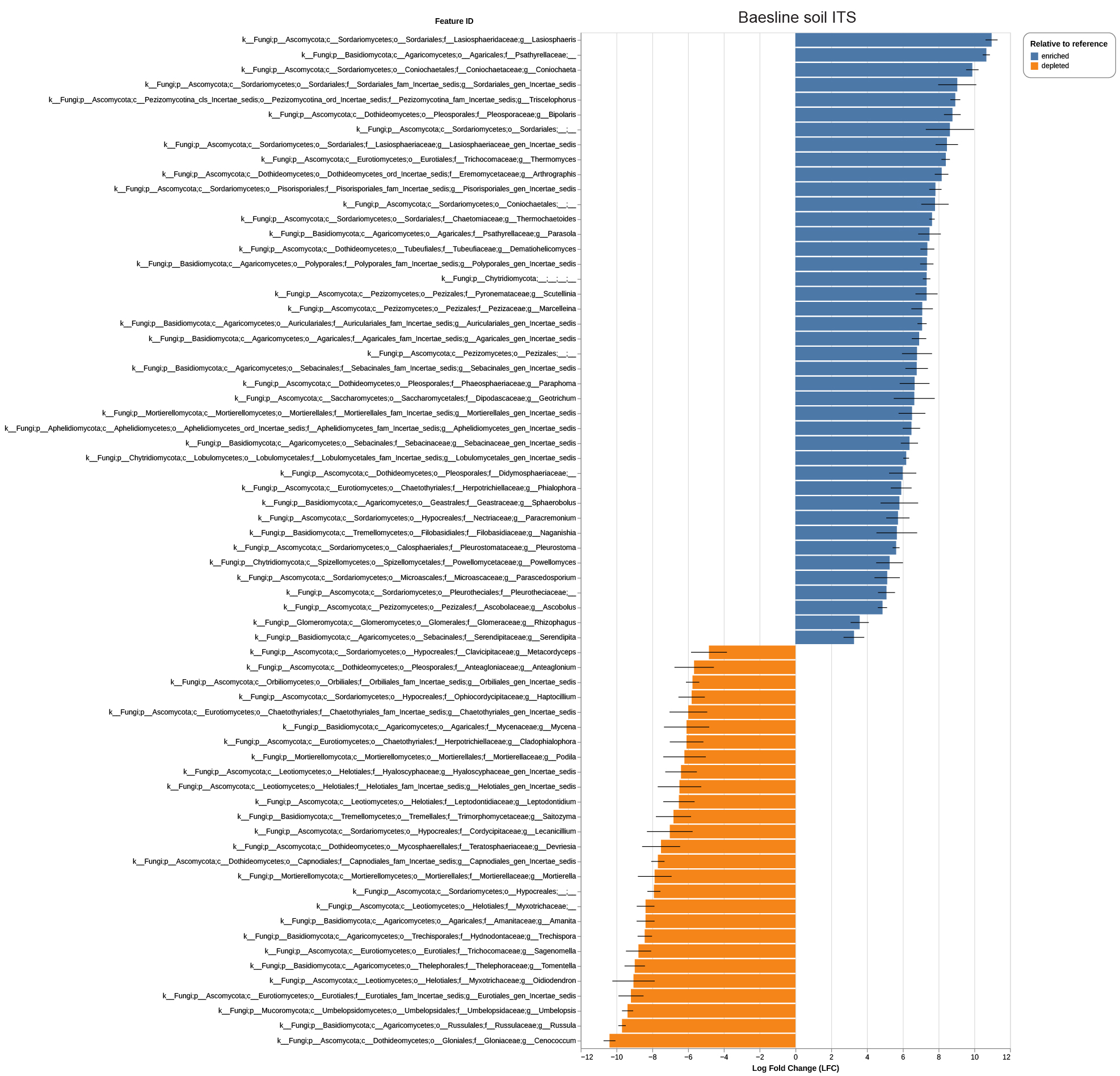

### FigS11

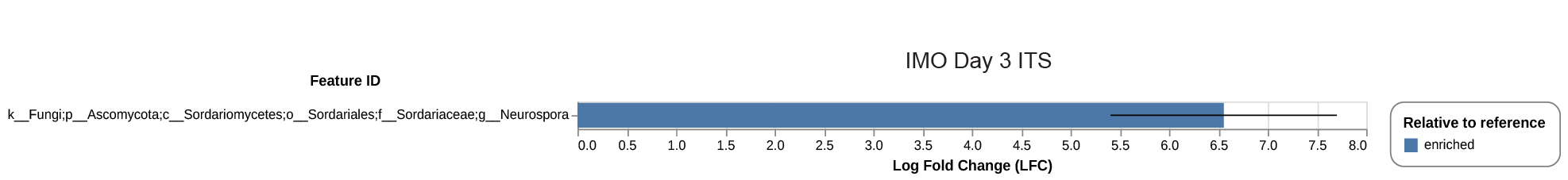

### FigS12

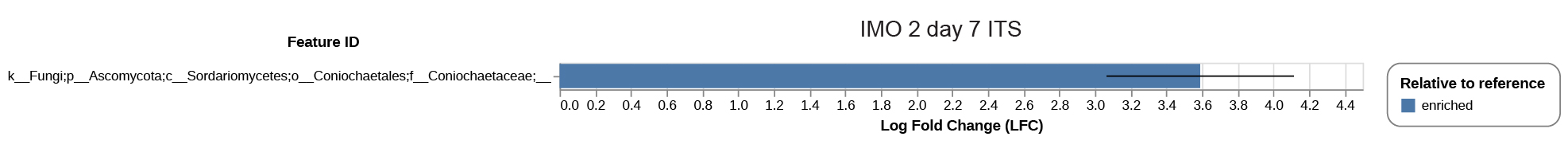

### FigS13

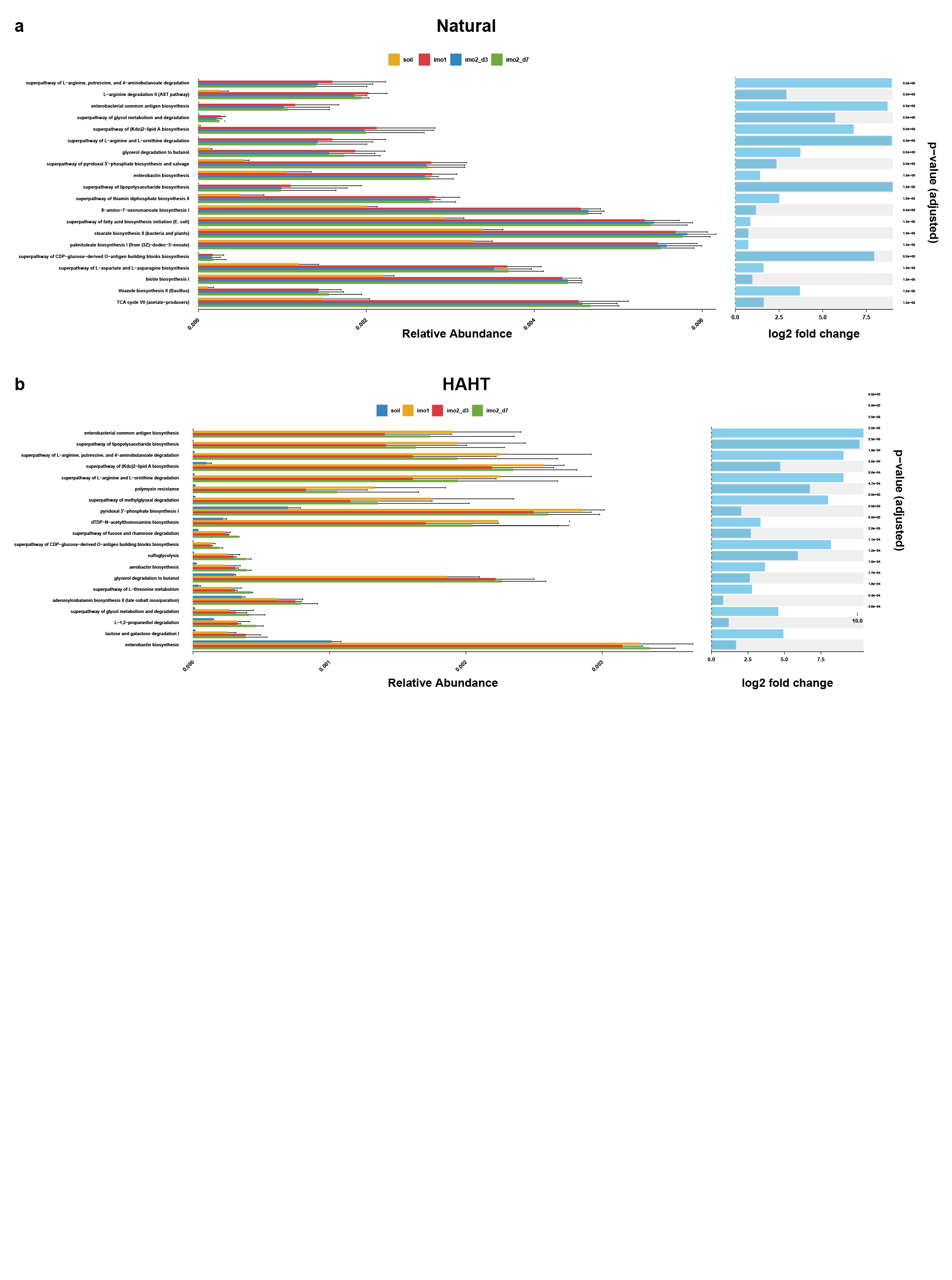
